# The specificity of sperm-mediated paternal effects in threespined sticklebacks

**DOI:** 10.1101/2020.06.04.135152

**Authors:** Eunice Chen, Christian Zielinski, Jack Deno, Raiza Singh, Alison M Bell, Jennifer K Hellmann

**Affiliations:** Department of Evolution, Ecology and Behavior, School of Integrative Biology, University of Illinois Urbana-Champaign, Urbana, Illinois, USA, 61801; Carl R. Woese Institute for Genomic Biology, University of Illinois Urbana-Champaign, Urbana, Illinois, USA, 61801; Program in Ecology, Evolution and Conservation, University of Illinois Urbana-Champaign, Urbana, Illinois, USA, 61801

**Keywords:** phenotypic plasticity, *Gasterosteus aculeatus*, nongenetic inheritance, predation risk, transgenerational plasticity, predator recognition

## Abstract

Parental effects can help offspring cope with challenging environments, but whether these effects are unique to specific environmental conditions is largely unknown. Parental effects may evolve via a core pathway that generally prepares offspring for risky environments or could be stimuli-specific, with offspring developing phenotypes that are tailored to specific environmental challenges. We exposed threespined sticklebacks (*Gasterosteus aculeatus*) fathers to a potentially threatening stimulus (net) versus native predator (sculpin). Offspring of sculpin-exposed fathers were more responsive (greater change in activity) to a simulated predator attack, while offspring of net-exposed fathers were less responsive (lower plasma cortisol and fewer antipredator behaviors). To evaluate offspring response to native and non-native stimuli, we sequentially exposed offspring of net-exposed, sculpin-exposed or control fathers to a net, native sculpin model, or non-native trout model. Paternal treatment did not influence offspring response to stimuli; instead, offspring were more responsive to the native sculpin predator compared to nets or non-native trout predator. Collectively, we demonstrate that sperm-mediated paternal effects in response to different, potentially stressful stimuli result in distinct offspring phenotypes. This specificity may be key to understanding the evolution of adaptive parental effects and how parents prime offspring for encountering both evolved and novel environmental stimuli.

## Background

Transgenerational plasticity (environmental parental effects) occurs when the environment experienced by one generation influences the phenotypes of future generations. Transgenerational plasticity may have evolved as a mechanism to help organisms to cope with changing environments and can have adaptive consequences for offspring (reviewed in [1–3]). Despite growing evidence for parental effects in response to environmental challenges such as predation risk, the ways in which parental effects prime offspring for specific environmental conditions is largely unknown.

On one hand, there may one conserved pathway by which the parental environment alters offspring phenotypes, such that different environmental conditions experienced by parents (e.g., low food availability, drought, predation) have the same intergenerational consequences: parents produce offspring with traits that help them cope with low quality environments (e.g., dispersal, altered stress responsivity). For example, female bluebirds adjust sex-biased laying order to shift offspring dispersal in response to low nest cavity abundance, cold temperature, or reduced kin proximity [4]. This generalized response could be adaptive by allowing parents to convey a message of environmental risk, even if they lack certain information about environmental cues (e.g. do not have to evaluate if high predation is due to sit-and-wait predators or active foragers) or encounter a novel stressor (e.g., a non-native predator). Alternatively, it could arise because low quality environments generally increase parental stress and/or reduce body condition [5–7]. On the other hand, parental effects may be highly specific, such that different parental conditions have very different intergenerational effects. For example, maternal social instability stress in mice increased anxiety predominantly in female offspring, while restraint stress reduced anxiety in offspring of both sexes [8]. If parental cues are specific, transgenerational cues may induce tailored changes in offspring phenotypes (e.g., altered metabolism vs increased antipredator behavior) that are adaptive for coping with specific environmental challenges (e.g., food instability vs. predation).

To investigate the specificity of parental effects, we compared the intergenerational effects of exposing fathers to a native predator (model sculpin *Cottus asper*) or to an artificial stressor (nets). There is abundant evidence that parental exposure to predation risk can induce offspring traits related to predator defense, such as protective “helmets” in *Daphnia* [9], improved antipredator behavior in crickets [10], and altered activity and exploration under high risk conditions in sticklebacks [11]. Artificial stimuli, such as nets, foot shocks or confinement, are often used as proxy for a predator or to induce acute stress [12–14] and can induce intergenerational effects [15, 16]. Given that individuals have not encountered these stimuli in their evolutionary history, it is unclear how individuals perceive them and if they induce the same intergenerational response as a natural predator. We then examined offspring of control fathers, sculpin-exposed fathers, and net-exposed fathers for a variety of antipredator traits, including antipredator behaviors, acute (cortisol) stress, and body size. We specifically tested the hypothesis that predator-induced paternal effects are specific, i.e. that offspring of predator-exposed fathers differ from offspring of both control and net-exposed fathers.

We paired this initial investigation of the specificity of paternal effects with a second experiment to understand if stickleback behave differently in response to a net compared to a model predator, which would suggest that they perceive nets and predators differently. For instance, individuals might respond less strongly or quickly show a diminished response to less threatening stimulus [17–19]. We sequentially exposed offspring of control, net-exposed, and sculpin-exposed fathers to either a net, model sculpin, or model trout predator (another stickleback predator that is not native to our population). This experiment also allowed us to determine whether parental exposure to a stimulus primes offspring response to that stimulus, e.g. if offspring of sculpin-exposed fathers respond more strongly to sculpin while offspring of net-exposed fathers respond more strongly to respond to a net.

## Method

### Housing conditions

Adult threespined sticklebacks were collected from Putah Creek, a freshwater stream in Davis, California, in August 2017. This population has piscivorous predators, including the prickly sculpin (*Cottus asper*). In October-November 2017, males were transferred to individual 26.5L nesting tanks; once they built nests, males were randomly assigned to a treatment in which they were either chased for 30 seconds every other day for a twelve-day period (6 times total) with a net (10 by 12.5cm, green) or a clay model sculpin (21cm length) or left undisturbed for an equivalent amount of time. Previous experiments have shown that stickleback respond most strongly to visual cues of predation [20] and model predators elicit anti-predator behaviors when brought into close proximity to sticklebacks [21]. Males were not transferred between tanks using nets before the experiment began to ensure that fathers were not habituated to the presence of a net; multiple nets of the same color and size were used to chase fathers.

The day after the last exposure, we removed the male, extracted his testes, and used his sperm to fertilize eggs of a wild-caught, unexposed female. We placed fertilized eggs in a cup with mesh bottom above a bubbler and monitored for mold before hatching. Stickleback males produce sperm in the beginning of the breeding season; thus, paternal effects mediated via sperm in this experiment are likely due to modifications to already mature sperm [22]. Because mothers and fathers did not interact with each other prior to fertilization or with their offspring post-fertilization, we could isolate paternal effects mediated via sperm and eliminate other potential sources of paternal effects, such as mate choice and differential allocation mediated via parental care [23]. We generated 17 clutches: n = 6 clutches of control fathers, n = 6 clutches of net-exposed fathers, n = 5 clutches of sculpin-exposed fathers. In accordance with a previous study on sperm-mediated paternal effects in this population [24], we found no detectable effect of paternal identity on any offspring traits whose mean value was significantly altered by paternal treatment (unpublished analyses), suggesting that genetic variation among clutches does not substantially alter these intergenerational effects.

### Part I: Transgenerational plasticity in response to an artificial stimuli (net) versus a native predator (sculpin)

#### Behavioral assays

When offspring were 5 months old (mean days post-hatching: 145.9 ± 0.64 s.e.; February-April 2018), we measured activity and antipredator behavior using methods described in Hellmann, Carlson [24] (5-13 offspring per clutch). The testing arena was a circular arena (150cm diameter) divided into eight sections on the perimeter with a middle circular section. We gently caught a fish from their home tank with a cup and transferred it to an opaque container in the middle section. After a three minute acclimation period, we removed the plug from the refuge, allowed the individual to emerge, and measured the number of sections visited for three minutes after emergence as a proxy for activity. If the fish did not emerge after 5 minutes, it was gently released from the refuge; whether fish emerged naturally or were released did not alter activity (Welch’s two sample t-test: t_180.12_=-0.22, p=0.83).

After the initial observation period, we simulated a sculpin predator attack by quickly moving a clay predator sculpin toward the section of the pool with the experimental fish. We measured two antipredator behaviors: whether this attack elicited evasive swimming behavior (binomial) and how long the fish spent frozen after the simulated predator attack (continuous). Once the fish resumed movement, we again measured the total number of sections visited for 3 minutes. We assayed n = 59 offspring of control fathers (n = 30 males, n = 29 females), n = 64 offspring of net-exposed fathers (n = 33 males, n = 31 females), and n = 60 offspring of sculpin-exposed fathers (n = 28 males, n = 32 females).

We left the fish in the assay arena for 15 minutes after the simulated predator attack in order to measure peak circulating plasma cortisol in response to the predator attack [25]. We then netted the fish from the arena and quickly weighed and measured it (standard length: from the tip of the nose to the base of the caudal fin). We euthanized the fish in MS-222, cut off the tail, and drew blood from the tail of the fish using a heparinized microhematocrit tube. We centrifuged blood to separate the plasma (StatSpin CritSpin Microhemocrit centrifuge) and immediately froze the plasma at −80 °C for enzyme-linked immunosorbent assay (ELISA; see supplementary material). We visually sexed offspring when possible, as many fish had underdeveloped and non-reproductively mature gonads; we confirmed the accuracy of this method and sexed the remainder of the fish using a genetic marker, per the methods of Peichel, Ross [26]. Due to an insufficient amount of blood drawn from some offspring, our final sample size analyzed was n = 38 offspring of control fathers, n = 27 offspring of net-exposed fathers, and n = 31 offspring from sculpin exposed fathers.

#### Statistical analysis

For all traits, we tested for differences in variance due to paternal treatment using Fligner-Kileen tests. We computed the difference in activity before versus after the attack (e.g. sections visited before - visited after the simulated predator attack). We used linear mixed effects models to test predictors of activity differences and cortisol, and generalized linear mixed models to test predictors of freezing behavior (negative binomial distribution) and evasive swimming (binomial distribution) (R package lme4 [27]). The models all included fixed effects of paternal treatment (control, net-exposed, sculpin-exposed), offspring sex, and standard length; however, we used log-transformed tank density for the evasive swimming model instead because model comparisons showed tank density to significantly improve model fit compared to standard length (standard length and density are highly correlated; Spearman rank correlation, ρ = −0.51, p<0.001). We included a random effect of clutch identity for all models, and observer identity for all behavioral models. We removed 3 outliers from the cortisol dataset to normalize the residuals. We tested for interactions between fixed effects and removed all non-statistically significant interactions. For all models, we tested for differences among paternal treatment using Tukey’s HSD (package multcomp [28]).

### Part II: The behavioral response of offspring to different stimuli

#### Open field assays

In April-May 2018, we ran the behavior assay on a different set of offspring from the same clutches, using different stimuli to simulate a predator attack. Each individual was isolated in a 10L tank (L32 x W21 x H19 cm) for at least 24hrs prior to the first assay. Each individual was chased with three different stimuli in separate assays: 1) a net, 2) a model sculpin as above, and 3) a model trout (a stickleback predator that is not native to the parents’ population; 19.7cm length). Assays were conducted in a random order, each 2-3 days apart; due to some experimental issues, not all fish received all three assays. After the last assay, fish were euthanized with MS-222, weighed, measured, and sexed via visual inspection of non-reproductively mature gonads. We conducted a total of n=131 assays: n = 16 offspring of control fathers (n = 16 sculpin assays, n=15 net assays, n=14 trout assays), n = 16 offspring of net-exposed fathers (n = 15 sculpin assays, n=16 net assays, n=12 trout assays), and n = 15 offspring of sculpin-exposed fathers (n = 15 sculpin assays, n=15 net assays, n=13 trout assays).

#### Statistical analysis

We reran the same behavioral models as above, with additional fixed effects of assay treatment (net, sculpin, trout) and assay number (first, second, or third assay). We included random effects of observer identity and fish ID nested with clutch. We tested for interactions between fixed effects and removed all non-statistically significant interactions. We removed one extremely low outlier from the activity dataset.

## Results

### Part I: Transgenerational plasticity in response to an artificial stimuli (net) versus a native predator (sculpin)

We observed different inter-generational effects of chasing fathers with either an artificial stimuli (net) or a native predator (sculpin).

#### Offspring behavior

Paternal treatment significantly altered offspring activity in response to a simulated predator attack (Table 1): offspring of sculpin-exposed fathers decreased activity more in response to the simulated predator attack compared to offspring of control fathers (Tukey’s HSD: z=1.93, p=0.13; Figure 1) while offspring of net-exposed fathers did not decrease activity as much as offspring of control fathers (z=-1.15, p=0.48; significant difference between offspring of net and sculpin-exposed fathers: z=3.19, p=0.004). Further, although we did not detect an overall effect of paternal treatment on offspring antipredator behavior (evasive swimming) (Table 1; Figure 2A), pairwise comparisons show that offspring of net-exposed fathers were significantly less likely to perform evasive swimming behavior compared to offspring of control fathers (z=-2.40, p=0.04). Offspring of sculpin-exposed fathers did not differ from offspring of control fathers (z=-1.45, p=0.32). Offspring freezing behavior was not detectably altered by paternal treatment (Table 1).

**Figure 1:**
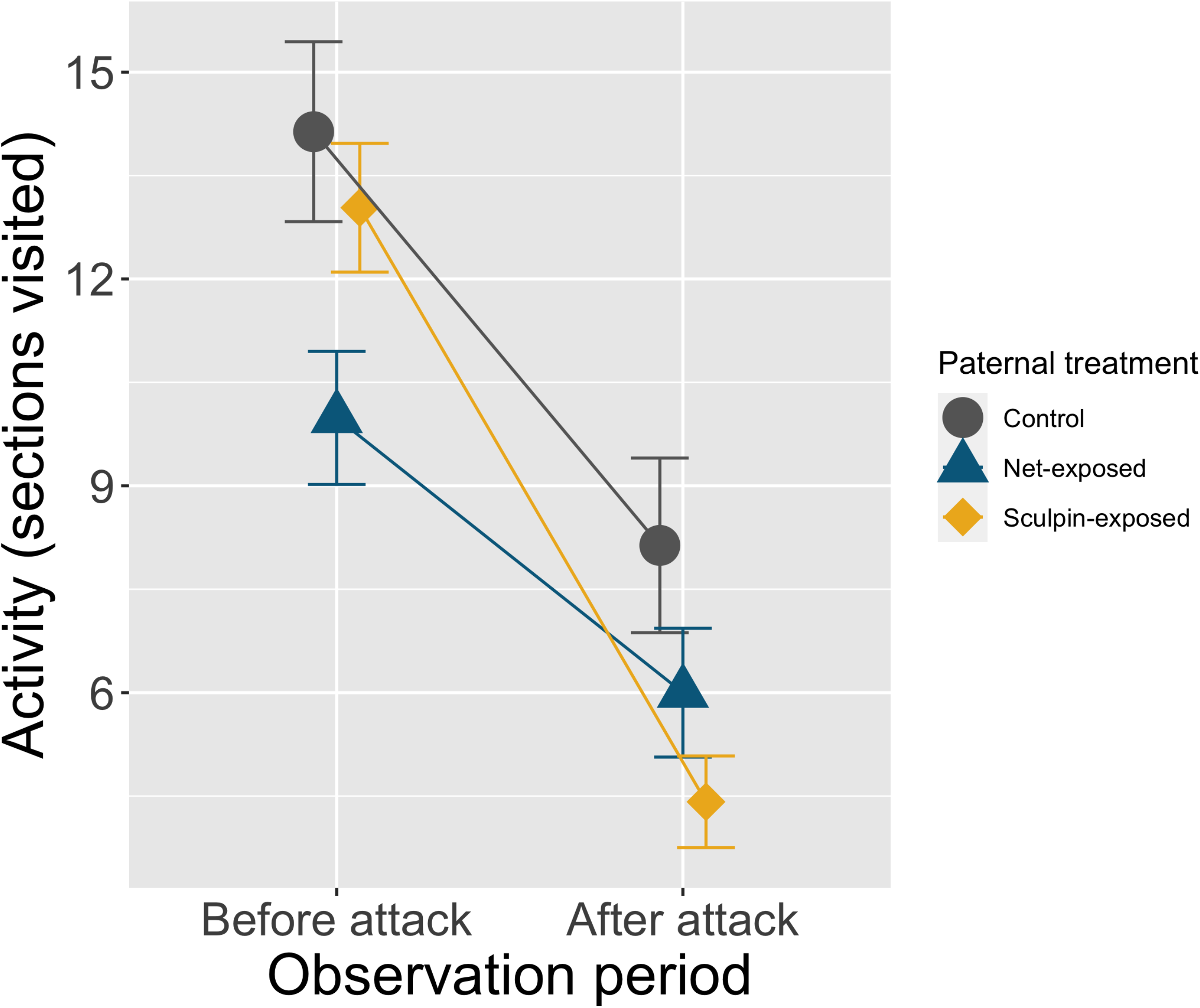
Relative to offspring of net-exposed fathers (blue), offspring of sculpin-exposed fathers (yellow) had a significantly greater reduction in activity in response to a simulated model sculpin attack in the open field assay (mean ± s.e.).

**Figure 2:**
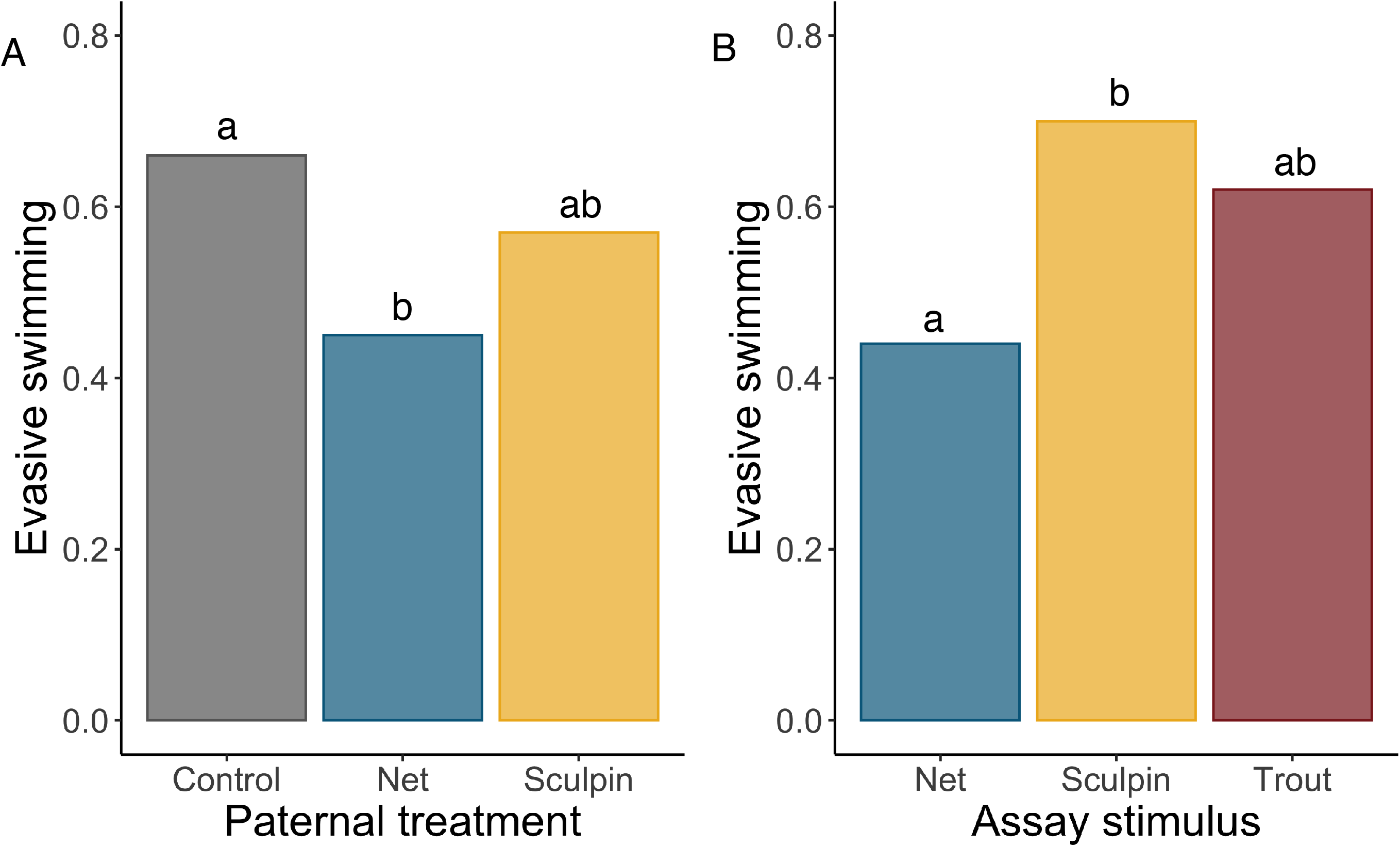
A) In Part I, paternal treatment (control, net-exposed, sculpin-exposed) significantly altered evasive swimming behavior of offspring in response to a simulated model sculpin attack in the open field assay. Shown are the proportion of individuals who performed this behavior (binomial response). B) In Part II, assay stimulus (net, sculpin predator, trout predator) significantly influenced whether offspring displayed evasive swimming behavior in response to a simulated attack in the open field assay. Letters indicate significant differences among treatment groups (Tukey’s HSD).

**Table 1:**
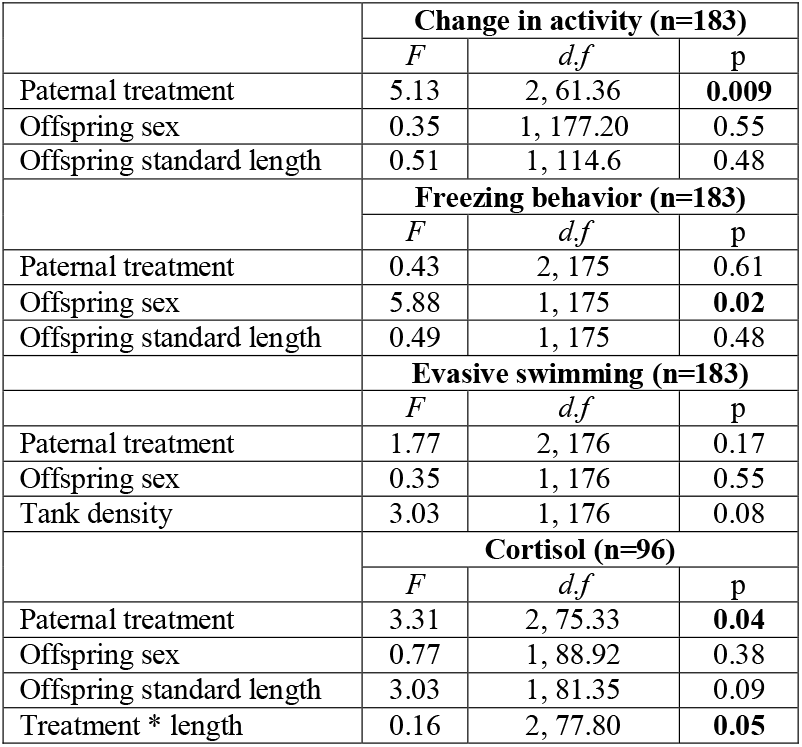
We used general linear mixed models to understand how paternal treatment altered offspring activity before versus after the simulated predator attack (higher values indicate a greater reduction in activity), freezing behavior, evasive swimming, and offspring cortisol responses at 5 months. All individuals underwent one open field assay, in which they were chased with a model predator sculpin.

|  | **Change in activity (n=183)** |  |  |
| --- | --- | --- | --- |
|  | *F* | *d.f* | p |
| Paternal treatment | 5.13 | 2, 61.36 | **0.009** |
| Offspring sex | 0.35 | 1, 177.20 | 0.55 |
| Offspring standard length | 0.51 | 1, 114.6 | 0.48 |
|  | **Freezing behavior (n=183)** |  |  |
|  | *F* | *d.f* | p |
| Paternal treatment | 0.43 | 2, 175 | 0.61 |
| Offspring sex | 5.88 | 1, 175 | **0.02** |
| Offspring standard length | 0.49 | 1, 175 | 0.48 |
|  | **Evasive swimming (n=183)** |  |  |
|  | *F* | *d.f* | p |
| Paternal treatment | 1.77 | 2, 176 | 0.17 |
| Offspring sex | 0.35 | 1, 176 | 0.55 |
| Tank density | 3.03 | 1, 176 | 0.08 |
|  | **Cortisol (n=96)** |  |  |
|  | *F* | *d.f* | p |
| Paternal treatment | 3.31 | 2, 75.33 | **0.04** |
| Offspring sex | 0.77 | 1, 88.92 | 0.38 |
| Offspring standard length | 3.03 | 1, 81.35 | 0.09 |
| Treatment * length | 0.16 | 2, 77.80 | **0.05** |

We did not detect an effect of offspring sex on activity or evasive swimming behavior, although males spent less time frozen than females, and we did not detect an effect of offspring length on activity or freezing behavior (Table 1). Tank density tended to increase the likelihood that offspring performed evasive swimming behaviors (Table 1). We did not detect differences in variance on change in activity (Fligner-Killeen test: χ^2^ = 1.42, p = 0.49) or freezing behavior (χ^2^ = 0.53, p = 0.77) between paternal treatments.

#### Offspring cortisol

Offspring of net-exposed fathers had lower cortisol following exposure to the model sculpin relative to offspring of control fathers (Tukey’s HSD: z=-2.50, p=0.03, Table 1; Figure 3); we did not detect a difference between offspring of control and sculpin-exposed fathers (z=-0.23, p=0.97). Offspring body size interacted with paternal treatment to influence cortisol (Table 1): large offspring of net-exposed fathers had higher cortisol (F_1, 24.00_=8.08, p=0.009), but this pattern was not apparent for offspring of control fathers (F_1, 21.30_=0.04, p=0.84) or sculpin-exposed fathers (F_1, 29_=0.01, p=0.92). We did not detect sex differences in cortisol (Table 1), and there was no evidence that paternal treatments differed in variance (χ^2^ = 0.26, p = 0.88).

**Figure 3:**
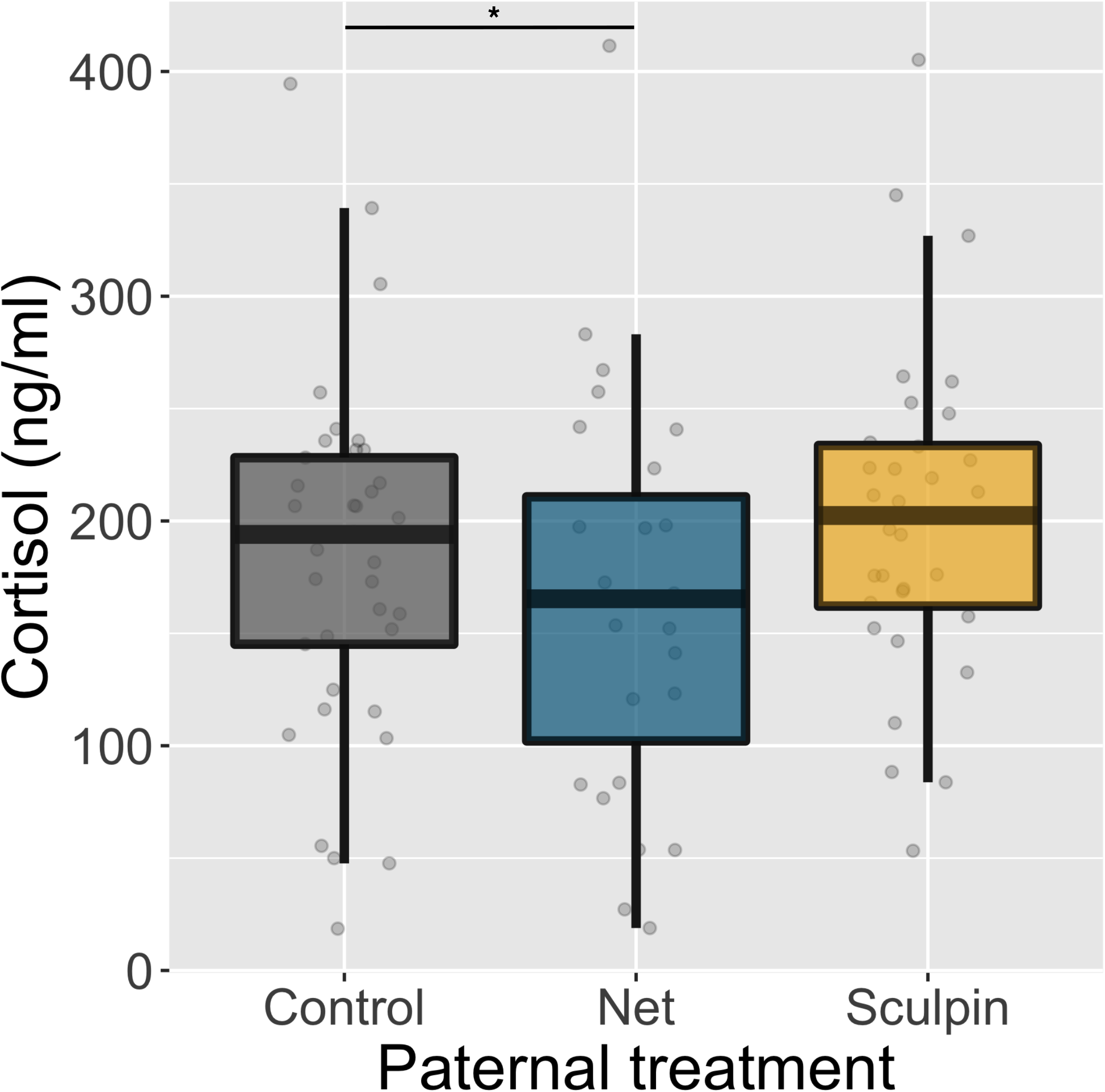
Relative to offspring of control fathers (grey), offspring of net-exposed fathers (blue) had significantly lower circulating cortisol (ng/ml) 15 minutes after a simulated model sculpin attack in the open field assay (interquartile range with median). These patterns were not present for offspring of sculpin-exposed fathers (yellow). Grey circles represent individual data points.

#### Offspring body size

We did not detect paternal effects on mean offspring standard length or mass (supplementary material). However, paternal treatments differed in the variance of offspring standard length and mass (Fligner-Killeen test; mass: χ^2^ = 14.50, p = 0.002; SL: χ^2^ = 11.89, p = 0.008). Specifically, offspring of net-exposed fathers were more variable in length and mass than offspring of sculpin-exposed fathers (SL: χ^2^ = 5.17, p = 0.02; mass: χ^2^ = 12.85, p <0.001); offspring of net-exposed fathers were also more variable in length than offspring of control fathers (χ^2^ = 6.44, p = 0.01). Neither of these differences in variance could be attributed to higher within-clutch coefficients of variation (SL: F_2,16_=0.18, p=0.68; mass: F_2,16_=0.28, p=0.84), suggesting that they are due to variation among fathers.

### Part II: Offspring response to different stimuli

We ran a second set of experiments in which each offspring was sequentially exposed to an artificial stimulus (net), a native model sculpin predator, and a non-native model trout predator. We observed different behavioral responses depending on which stimulus offspring encountered (Table 2: evasive swimming). Specifically, sticklebacks were more likely to perform evasive swimming behaviors when they encountered a sculpin compared to a net (Z=2.54, p=0.03), but we did not detect a difference between stickleback’s response to a sculpin versus a trout (Z=1.06, p=0.54; Figure 2B, Table 2).

**Table 2:** We used general linear mixed models to understand offspring’s response to different stimuli (n=131 assays). We tested the effects of paternal treatment (control, net-exposed, and sculpin-exposed) and assay stimulus (sculpin, net, trout) on offspring activity behavior (higher values indicate a greater reduction in activity after the predator attack), freezing behavior, and evasive swimming (binomial). Individuals underwent three open field assays, and were chased with each of the three assay stimuli in a randomized order.

|  | **Change in activity** |  |  |
| --- | --- | --- | --- |
|  | *F* | *df* | p |
| Paternal treatment | 0.70 | 2, 7.84 | 0.52 |
| Assay stimulus | 3.90 | 2, 116.54 | **0.02** |
| Assay number | 28.59 | 1, 82.34 | **<0.001** |
| Offspring sex | 0.16 | 1, 41.71 | 0.69 |
| Offspring standard length | 0.81 | 1, 16.63 | 0.38 |
| Assay stimulus * assay number | 3.30 | 2, 115.10 | **0.04** |
|  | **Freezing behavior** |  |  |
|  | *F* | *df* | p |
| Paternal treatment | 1.13 | 2, 118 | 0.33 |
| Assay stimulus | 0.56 | 2, 118 | 0.57 |
| Offspring sex | 0.09 | 1, 118 | 0.76 |
| Assay number | 12.39 | 1, 118 | **<0.001** |
| Offspring standard length | 0.70 | 1, 118 | 0.40 |
|  | **Evasive swimming** |  |  |
|  | *F* | *df* | p |
| Paternal treatment | 2.53 | 2, 120 | 0.08 |
| Assay stimulus | 4.08 | 2, 120 | **0.02** |
| Offspring sex | 0.12 | 1, 120 | 0.72 |
| Assay number | 0.10 | 1, 120 | 0.75 |
| Tank density | 0.97 | 1, 120 | 0.33 |

Because individuals were repeatedly run through the behavioral assay, we could examine how behavior changed with repeated testing in the behavioral assay. We detected effects of repeated testing: regardless of the assay stimulus, offspring spent more time frozen after the simulated attack in the first assay compared to later assays (Table 2). However, there were more pronounced effects of repeated testing in response to some stimuli compared to others (assay stimulus by assay number interaction for activity; Table 2; Figure 4). For example, stickleback showed a greater reduction in activity after being chased by the net in their first assay compared to later assays (F_1, 39_=13.33, p<0.001). The same pattern was evident for trout (F_1, 26.37_=16.23, p<0.001). In contrast, when individuals were chased with a sculpin, they showed similar reductions in activity in the first assay compared to later assays (F_1_, 39.97=2.69, p=0.11). This suggests that stickleback are less responsive to nets and non-native trout predators following experience in the behavioral assay, but remain responsive to native sculpin predators over time.

**Figure 4:**
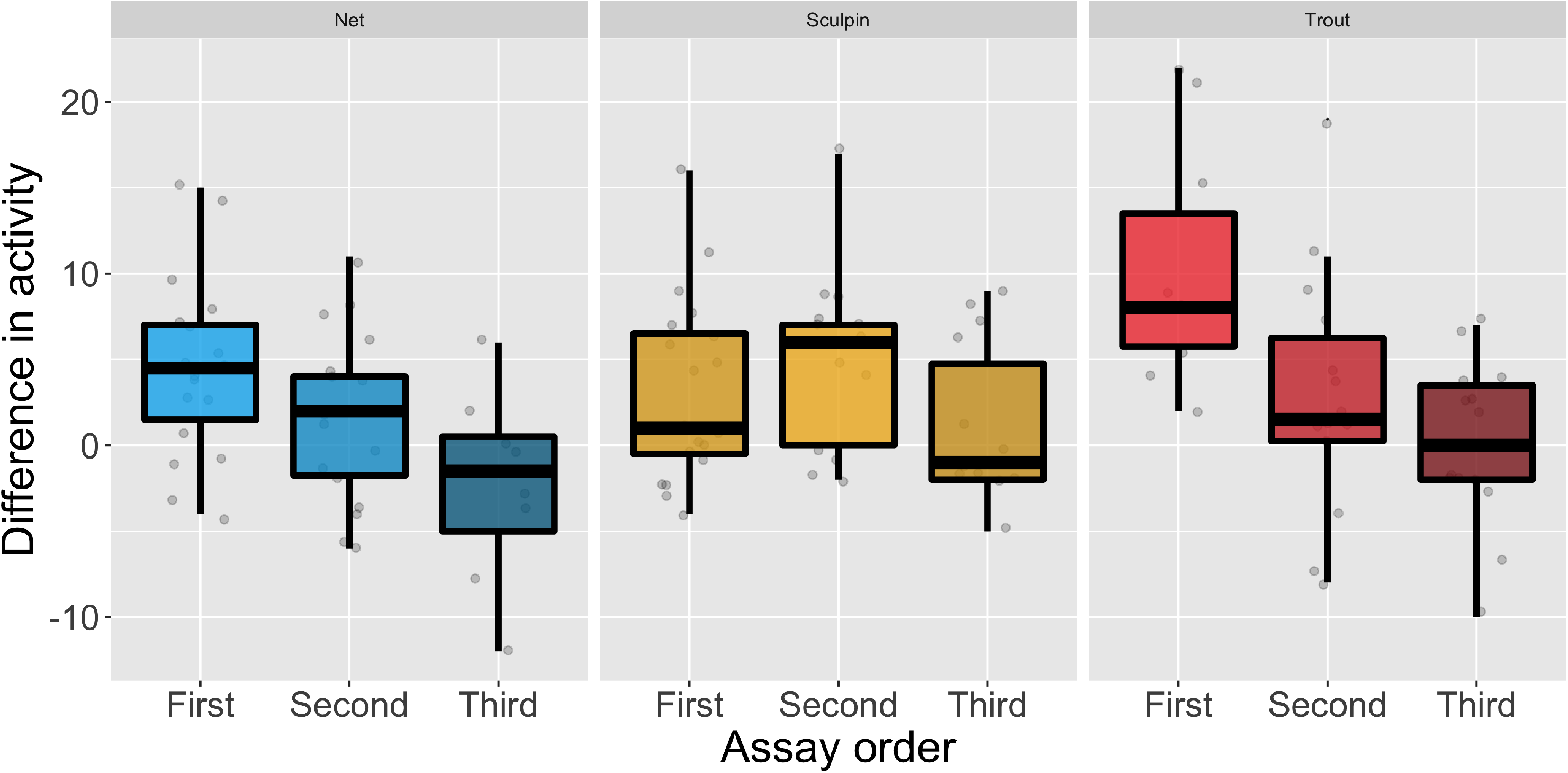
In Part II, offspring who encountered a non-native trout predator or a net show a smaller decrease in activity in response to the simulated predator attack in later assays compared to the first assay (interquartile range with median). However, this trend was not present for offspring who were exposed to a native sculpin predator in the assay. Grey circles represent individual data points. See Supplementary Figure 1 for plot of raw data.

Because we measured the behavior of offspring of net-exposed versus sculpin-exposed fathers, we looked for evidence that parental exposure to a given stimulus primes offspring to respond to that stimulus (e.g., if offspring of net-exposed fathers are primed to respond to nets). We did not detect evidence for paternal priming, as evidenced by non-statistically significant interactions between parental treatment and assay stimulus (Table 2). The direction and magnitude of the paternal effects in Part I and Part II are similar (see supplementary material); the failure to detect a significant effect of paternal treatment in Part II likely reflects its smaller sample size.

## Discussion

Here, we sought to investigate the specificity of transgenerational plasticity by examining how offspring phenotypes varied when stickleback fathers were exposed to a general, non-ecologically relevant stimulus (net) versus a native predator (sculpin). After a simulated predatory sculpin attack, offspring of sculpin-exposed fathers showed a greater reduction in activity compared to offspring of net-exposed fathers. On the other hand, offspring of net-exposed fathers were less likely to perform antipredator behaviors (evasive swimming) and had significantly lower circulating cortisol in response to the simulated sculpin attack than offspring of control fathers, but these differences were not apparent for offspring of sculpin-exposed fathers. This is consistent with previous findings that paternal exposure to non-predation stress, transmitted to offspring via epigenetic changes to sperm, reduces HPA stress axis responsivity and lowers offspring anxiety [29, 30]. Collectively, these results demonstrate that paternal experience with different stimuli (net versus a predator) alter offspring phenotypes in distinctive ways.

Consistent with other studies [10, 31, 32], offspring of predator-exposed fathers were more responsive to simulated predation risk, in that they showed a greater reduction in activity in response to risk compared to offspring of unexposed fathers. Given that prey often decrease activity levels under high predation risk [33, 34], this suggests that paternal experience with a native predator heightens antipredator behavior. In contrast, paternal exposure to a non-ecologically relevant stimulus (nets) actually decreased antipredator behavior in response to a simulated sculpin attack. It is possible that nets and predators elicit different responses in fathers, which could explain why paternal effects in response to a net versus a predator were different. Consistent with this explanation, we found that stickleback behave differently toward nets and predators in Part II, which suggest that they perceive them differently.

If different parental stimuli result in different offspring responses, then it is possible that parents might be able to convey specific information about the stimulus that they encountered. For example, offspring may be primed to respond to the same stimulus that their father encountered (i.e., offspring of net-exposed fathers are more responsive to nets while offspring of sculpin-exposed fathers are more response to sculpin). We did not detect statistical evidence for this possibility; instead, we found that stickleback were generally more responsive to a native model sculpin predator compared to a net or non-native model trout. Namely, offspring were more likely to perform antipredator behaviors when they encountered a sculpin compared to a net; further, across the trials, offspring became less responsive to nets and non-native trout predators, but not to native sculpin predators. This is consistent with previous findings that individuals adjust the intensity of antipredator behavior after assessing the nature and overall threat of predation risk and that less threatening stimuli elicit an attenuated response over time [17–19].

Given that the animals used in Part II were previously predator-naïve, our results suggest that sticklebacks have innate predator recognition of native predators, even in the absence of olfactory or chemical predation cues. A number of fishes show innate predator recognition [35–37]. Although there is some evidence that learned predator avoidance is important in sticklebacks [38, 39], innate predator recognition may also occur: a previous study in sticklebacks also found that overhead fright response to birds is independent of predation experience [40]. This innate recognition of sculpin may arise from predator coloration or functional morphology (e.g., shape) [41, 42]. Indeed, our results do suggest that predator shape or coloration may be important for generalizing across different predators: stickleback that encountered a trout showed intermediate levels of antipredator (evasive swimming) behavior compared to stickleback who encountered a sculpin or net. This suggests that sticklebacks may be able to partially generalize predation risk to a non-native predator.

In addition to changes in mean offspring traits, we also found higher variance in standard length and body mass in offspring of net-exposed fathers, but not in offspring of sculpin-exposed fathers. Given that we found no differences in within-clutch coefficients of variation between treatment groups, our data suggest that higher variance cannot be attributed to ‘bet-hedging’, where each father produces offspring with a wide range of different phenotypes [43]. Instead, these shifts in variance likely result from differences among fathers in how they responded to the net treatment. This is consistent with previous results in sticklebacks that found that fathers who encountered a novel predator showed a more variable changes in paternal care behaviors relative to fathers who encountered a native predator, who showed consistent directional changes in behavior [44].

In conclusion, here we show that prefertilization paternal exposure to both an artificial stimulus (net) and a native predator induce changes in offspring phenotypes; however, each stimulus induced different changes in offspring phenotypes, demonstrating that sperm-mediated paternal effects can be highly specific to the stimulus fathers encounter. Our study suggests that, while non-ecologically relevant stimuli elicit effects in intergenerational studies, caution should be used when trying to extrapolate those findings to understand intergenerational effects in response to evolved stimuli such as predators. Further, these results are consistent with the idea that different parental stimuli do not activate a core, conserved pathway, although there may be common offspring traits, such as dispersal [4] or body size [45–47], that are altered by cues indicating low quality or risky environments. Instead, our results are consistent with the hypothesis that that different sensory neurons or neuroanatomical pathways are activated in response to each stimulus and somehow alter sperm content (e.g., small RNAs [48–50]) in distinct ways, similar to the ways in which paternal exposure to different odors can induce specific aversions in offspring [15]. Further investigation into the underlying mechanisms, as well as the fitness consequences of each exposure with respect to survival against predators, would be exciting avenues for future work.

## Acknowledgements and funding

Thank you to the Bell lab for comments on previous versions of this manuscript and to Ryan Earley for help with the hormone assays. This work was supported by the National Institutes of Health award number 2R01GM082937-06A1 to AMB and National Institutes of Health NRSA fellowship F32GM121033 to JKH. The authors have no conflicts of interest.

## Author contributions

In Part I, EC and CZ generated offspring, conducted behavioral assays and offspring sexing, and processed the cortisol samples. JD and RS conducted behavioral assays for Part II. JKH collected cortisol samples and conducted the statistical analyses. EC, CZ, and JKH designed the studies and drafted the manuscript. AMB provided space and equipment for the experiments, financial support, helped with the statistical analyses, and edited the manuscript with JKH. All authors gave final approval for publication and agree to be held accountable for the work performed therein.

