## Supplementary Material for "The specificity of sperm-mediated paternal effects in threespined sticklebacks"

***Supplementary methods: housing conditions.*** The parental generation was maintained on a summer photoperiod schedule (16 L : 8D) at 20° ± 1°C and fed ad libitum daily with a mix of frozen bloodworm (*Chironomus* spp.), brine shrimp (Artemia spp.) and Mysis shrimp. Males were in mixed sex stock tanks with gravel and plastic plants prior to nesting, and then transferred to individual 26.5L tanks (36L x 33W x 24H cm) with gravel, two plastic plants, a sandbox, a clay pot, and algae for nest building

Offspring were reared in 26.5L tanks (36L x 33W x 24H cm), with each clutch housed individually. Once hatched, offspring were fed newly hatched brine shrimp for two months and then frozen food (as above) for three months before starting the behavior assays. Offspring were switched to a winter light schedule (8 L: 16 D) within two months of hatching and were maintained on that schedule for all assays.

***Supplementary methods: plasma cortisol extraction***. To detect offspring’s cortisol response in response to the simulated predator attack, we followed the manufacture’s protocol (Enzo Life Sciences, Plymouth Meeting, PA, USA). All the samples were prepared in 1:1 steroid displacement reagent solution, in 1:120 dilution, and run in duplicate. Slopes of the standard curves and a serial dilution curve are parallel [1]. All samples with a coefficient of variation greater than 20% were excluded from further analysis. The intra-assay coefficients of variation (calculated from remaining samples) were all within acceptable range (3.4%, 8.1%, 6.7%, 7.4%, 4.2%, 5.9%). We ran common samples of pooled plasma on each plate (in quadruplicate as the first two and last two wells of each plate) to calculate the interassay coefficient of variation (18.4%).

***Supplementary results: mass and length analysis***

Because standard length and mass were heteroskedastic, we used MCMC generalized linear mixed models (R package MCMCglmm) with a weak prior on the variance (V=1, nu=0.002). We ran models for 200,000 iterations, with a burn-in of 3000 iterations, thin = 3, and Gaussian distributions. For the standard length model, we included fixed effects of parental treatment, offspring sex, days since hatched, and tank density. For the mass model, we included fixed effects of parental treatment, offspring sex, and standard length. For both models, we included a random effect of clutch identity.

We found no difference in length or mass between offspring of control fathers compared to offspring of net-exposed fathers (SL: 95% CI (-2.57, 3.15), p=0.80; mass: 95% CI (-0.06, 0.05), p=0.88) or sculpin-exposed fathers (SL: 95% CI (-3.57, 2.60), p=0.74; mass: 95% CI (-0.08, 0.03), p=0.41). Male and female offspring did not differ in length (95% CI (-0.01, 0.03), p=0.46) or mass (95% CI (-0.03, 0.01), p=0.48). Younger offspring (95% CI (0.01, 0.24), p=0.04) and offspring in higher density (95% CI (-0.28, -0.11), p<0.001) tanks were shorter. Heavier fish were also longer (95% CI (0.039, 0.045), p<0.001).

***Supplementary results: effects of paternal treatment in Part II***

In contrast to Part I, we found no overall effect of parental treatment on changes in offspring activity (Table 2). However, if we analyze only the first assay encountered by each individual (parallel to Part I, where each individual only underwent one assay instead of three), we do find the same general (albeit non-significant) paternal treatment effect (F_2, 39.38_=2.58, p=0.09): offspring of predator-exposed fathers showed a greater change in activity before and after the simulated predator attack compared to offspring of control fathers (t_39.15_=1.94, p=0.06), but this pattern was not true for offspring of net-exposed fathers (t_39.67_=-0.07, p=0.95). It is likely a weaker effect because Part II has a much smaller sample size than Part I (n=47 individuals as opposed to n=183).

As with Part I, we found no effect of parental treatment on offspring freezing behavior (Table 2), but we did find evidence that parental treatment altered evasive swimming behaviors: offspring with a net-exposed (Tukey’s HSD: Z= -2.39, p=0.04), but not a sculpin-exposed father (Z= -1.99, p=0.12), were less likely to perform evasive swimming behaviors than offspring of control fathers.

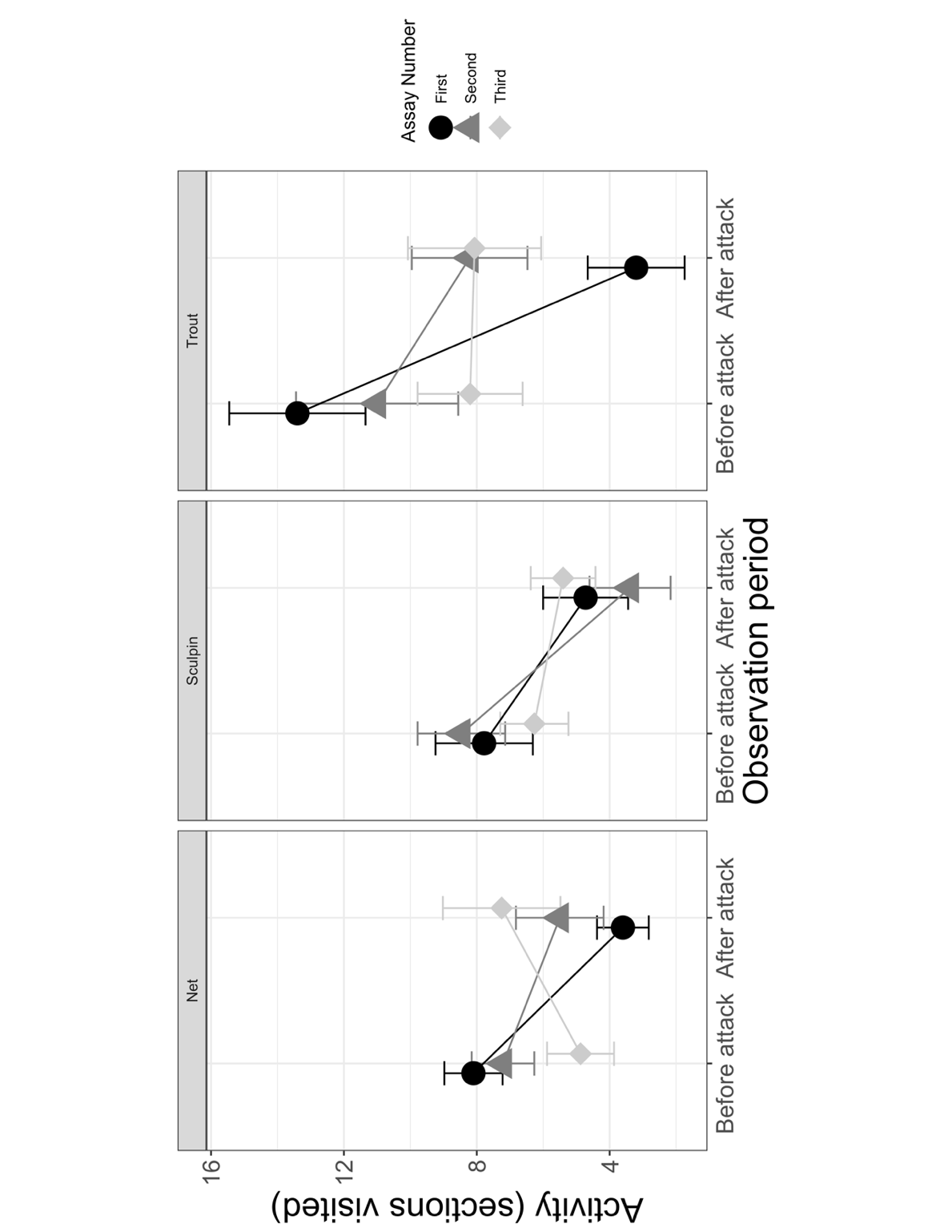

**Supplementary Figure 1:** In Part II, offspring who encountered a non-native model trout predator or a net show a reduced change in activity in later assays compared to the first assay (mean ± s.e.). However, this trend was not present for offspring who were exposed to a native model sculpin predator in the assay.
